# Alpha oscillations support attentional orienting while beta supports perceptual decision-making

**DOI:** 10.64898/2026.06.22.733411

**Authors:** Francesca M. Nannetti, Matias J. Ison, Mireia Torralba, Domenica Veniero

## Abstract

Visuospatial attention enables the selective allocation of cognitive resources to relevant stimuli. A well-established neural signature of attentional shifts is the lateralised modulation of occipito-parietal alpha power, with decreases over the hemisphere contralateral to the attended location and increases over the ipsilateral hemisphere. However, growing evidence suggests that multiple oscillatory mechanisms contribute to attentional deployment, including beta-band activity. A key unresolved question that remains is whether the same neural rhythms support the deployment of attention and the perceptual decisions that follow. Here, we recorded EEG in 26 participants (22 females) during covert visuospatial orienting and investigated how alpha- and beta-band dynamics relate to behavioural measures, namely perceptual sensitivity (*d*′) and decision criterion (*c*), and whether attended location could be preferentially decoded from alpha- or beta-band activity. We found that pre-target beta phase significantly predicted decision criterion at earlier pre-target intervals, whereas perceptual sensitivity was predicted closer to target onset, suggesting that beta is related to both sensory gain and the perceptual decision. In contrast, decoding analyses revealed that attended location was most strongly discriminable from alpha-band activity, as confirmed by time-frequency analysis of decoding accuracy. Together, these findings suggest a functional dissociation between oscillatory mechanisms supporting attentional orienting and perceptual decision-making. Whereas alpha-band activity primarily reflects the allocation of attention, beta-band dynamics predict trial-by-trial variability in perceptual decisions.

**Significance Statement:** Although alpha oscillations are commonly viewed as the principal neural mechanism underlying visuospatial attention, mounting evidence shows that such a complex function requires the involvement of other brain rhythms. Our findings show that different components of attention are associated with distinct brain rhythms. While alpha activity primarily reflects where attention is allocated, beta oscillations relate to how sensory information is transformed into perceptual decisions. These results suggest that attentional orienting and perceptual decision-making rely on dissociable neural processes, providing a comprehensive account of how brain oscillations support goal-directed behaviour.

## Introduction

Visuospatial attention enables the selective allocation of resources to behaviourally relevant information, allowing observers to prioritise stimuli at specific locations in space (Posner, 1980; Wahn and König, 2017). Decades of research have demonstrated that directing attention alters neural activity throughout the visual system and improves behavioural performance (Carrasco, 2015). Among the neural signatures associated with attentional orienting, modulation of alpha-band activity (8–12 Hz) over the occipito-parietal cortex has emerged as the canonical marker of visuospatial attention. Specifically, attention directed towards one side of space is accompanied by reduced alpha power in the contralateral hemisphere and increased alpha power ipsilaterally (Thut et al., 2006; Rihs et al., 2009), leading to the influential view that alpha oscillations constitute a core mechanism of top-down attentional control.

Despite the prominence of alpha-band accounts, accumulating evidence suggests that attentional control cannot be fully explained by alpha dynamics alone. Recent theoretical frameworks propose that attention emerges through rhythmic interactions among multiple oscillatory processes (Landau and Fries, 2012; Fiebelkorn et al., 2018; Helfrich et al., 2018; Fiebelkorn and Kastner, 2019). In particular, beta-band activity has been implicated in top-down signalling within frontoparietal networks (Bastos et al., 2015; Michalareas et al., 2016; Richter et al., 2017) and has been proposed to contribute to attentional sampling and stimulus processing (Fiebelkorn and Kastner, 2019; Di Dona and Ronconi, 2023). Consistent with this view, neural synchronisation in the beta range has been linked to the current focus of spatial attention (Siegel et al., 2008). Moreover, causal manipulations of the frontal eye fields (FEF) have been shown to induce phase resetting of occipital beta oscillations and corresponding fluctuations in visual performance (Veniero et al., 2021), while also modulating bottom-up signals in the gamma band (Trajkovic et al., 2025).

Together, these findings challenge the notion that attentional control can be explained by alpha-band dynamics alone and raise the possibility that distinct frequency bands contribute differentially to attentive behaviour.

A critical unresolved question concerns the functions supported by these rhythms. Most previous work has focused either on neural signatures of attentional orienting (e.g., Thut et al., 2006) or on perceptual decision-making in the absence of explicit attentional manipulation (e.g., Zhou et al., 2021). Consequently, it remains unclear whether alpha- and beta-band activity contribute similarly to the deployment of attention and to the perceptual decisions that follow. This distinction is important because attentional orienting and perceptual decision-making are theoretically separable processes. Attention determines where processing resources are allocated, whereas perceptual decisions concern how sensory evidence is evaluated and translated into behaviour.

Signal detection theory provides a framework for dissociating these components through measures of perceptual sensitivity (*d*′) and decision criterion (*c*). Studies examining spontaneous fluctuations in oscillatory activity have linked alpha-band dynamics to changes in criterion (Limbach and Corballis, 2016; Iemi et al., 2017; Sherman et al., 2016), perceptual sensitivity (Zhou et al., 2021), confidence (Samaha et al., 2017), and broader perceptual decision-making processes (Samaha et al., 2020). However, because these studies primarily investigated perceptual decision-making without attentional orienting, they provide limited insight into how oscillatory dynamics contribute to attentive behaviour. As a result, the relationship between oscillatory activity, attentional control, and perceptual decision-making remains unresolved.

Here, we tested whether attentional orienting and perceptual decision-making rely on common or dissociable oscillatory mechanisms. Using a sustained visuospatial attention paradigm, EEG recordings, signal detection theory, and multivariate decoding analyses, we examined how pre-target alpha- and beta-band activity relate to behavioural sensitivity, decision criterion, and attentional states. If alpha oscillations constitute a general mechanism of attentional control, they should predict both attentional orienting and behavioural outcomes. Alternatively, if distinct oscillatory processes support different components of attentive behaviour, alpha and beta activity should exhibit dissociable relationships with attentional state and perceptual decisions. By directly contrasting these possibilities, the present study aims to clarify the functional contributions of oscillatory dynamics to visual attention and behaviour.

### Materials and Methods Subjects

Forty healthy volunteers were recruited overall (32 female), all of whom had normal-to-corrected vision. All completed an initial behavioural session, and from this, 31 participants were invited back to complete an EEG session (25 female). Exclusion was due to insufficient performance on the behavioural task (for criteria, see below). An additional 5 participants (3 female) were further excluded due to insufficient performance on the EEG session (same criteria). The final sample size for analysis was 26 participants (*M* = 22.23 years, *SD* = 3; 22 females). The study was approved by the School of Psychology Ethics Committee at the University of Nottingham. All participants provided informed consent prior to the study and received monetary compensation for their time.

### Experimental Design

#### Main task

Participants performed a visuospatial attention task (Figure 1A) during which they covertly attended to a cued hemifield for an entire block, rather than receiving trial-by-trial cues. This choice was motivated by the observation that presenting a directional cue at the start of each trial may cause strong preferential alpha-band activity (Spadone et al., 2021). We therefore adopted a block-wise design to minimise cue-specific orienting responses and better isolate sustained pre-target attentional dynamics. To encourage participants to maintain covert spatial attention throughout each block, cue validity was set to 80%.

**Figure 1.**
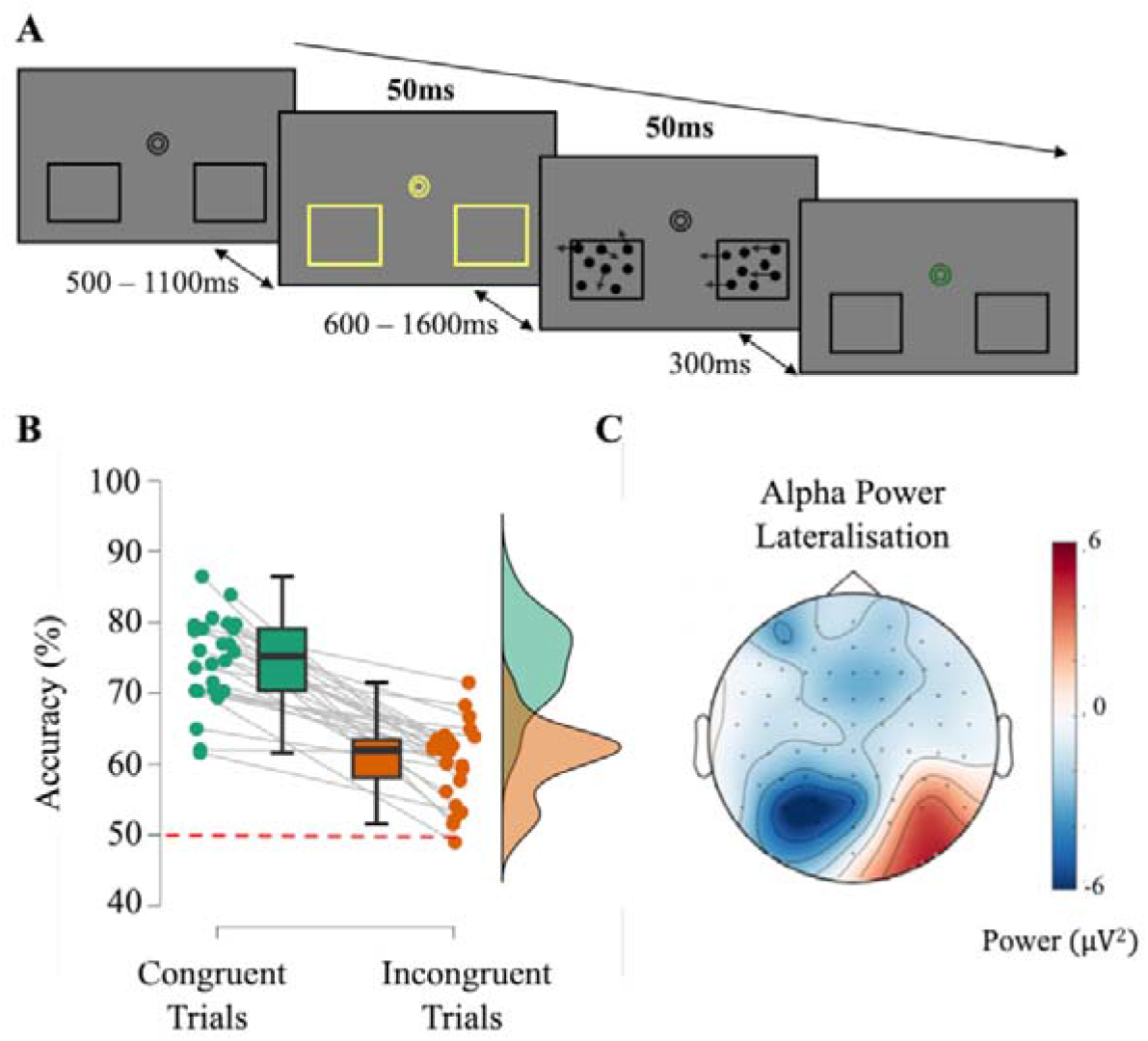
Behavioural task. A) Trial structure. After a jittered interval (500 – 1100ms), the fixation and placeholders changed colour for 50ms to indicate trial onset and therefore serving as a cue to initiate a covert shift of attention. Following a variable delay (600 – 1600ms), moving dot stimuli appeared in both placeholders for 50ms. One placeholder contained coherently moving dots (arrows shown for illustration), while the other contained randomly moving dots. In the example, the right placeholder contains coherent motion, and the left contains random motion. The fixation turned green 300ms after stimulus offset, prompting participants to report the direction of coherent motion. B) Behavioural performance. Boxplot of performance accuracy for congruent and incongruent trials. Chance performance (50%) is marked by a dashed red line. Each dot represents an individual participant; the box midline indicates the median, while whiskers represent the minimum and maximum data values. C) Attentional shift EEG signature. Grand average of rightward minus leftward orienting of attention in the alpha band (8-12 Hz) during the cue-target interval.

At the beginning of each block, participants were instructed with which hemifield would most likely contain the target (i.e., ‘Left Block’ or ‘Right Block’). There was a total of eight blocks, alternating between left and right attention conditions to reduce fatigue and maintain engagement.

A central bullseye and two placeholders were continuously presented on the screen. At a random time (500 – 1100ms), these briefly changed colour (yellow, isoluminant to the background), acting as a cue signalling trial onset and prompting participants to covertly shift their attention to the cued placeholder. After a variable interval (600 – 1600ms), two sets of moving dots (180 black dots, size: 5 degrees; field space: 5.8, speed: 0.04; lifetime: −1) appeared within each placeholder for 50ms. One set of these dots moved coherently and was, therefore, the target. The other moved randomly and was, therefore, the distractor. The order of congruent and incongruent trials was randomised within blocks, and target movement direction was equally likely to be either right or left for each trial. Participants discriminated the direction of coherently moving dots, responding after the fixation turned green (300ms following stimulus offset) using a keyboard with no time limit for response. Motion coherence was individually titrated to achieve ~75% accuracy at the cued location. For incongruent trials, coherence was increased by 10% to ensure that participants’ performance was above-chance at the uncued location, thereby encouraging participants to continue sampling the unattended location rather than strategically ignoring it. After each block, participants were visually presented with accuracy scores (%), followed by a self-paced rest period. Motion coherence was changed if participants were under- or overperforming: +5% if accuracy was < 70%, +10% if < 60%, −5% if > 80%, and −10% if > 90%, compensating for both practice and fatigue effects. The task was created and presented via PsychoPy (version 2021.2.3; Peirce et al., 2019).

### Procedure

Participants attended 2 sessions on separate days (behavioural and EEG). During both, they sat in a dim, sound-attenuated Faraday cage, placing their chin on a chinrest (~57cm from the screen) to minimise head movements and position a central line of sight. Eye position was monitored monocularly at 1000 Hz using an EyeLink 1000 (version 1.0.18; SR Research, Mississauga, Ontario, Canada). A standard 5-point calibration and validation was performed at the start of each task block (*N* = 4 calibrations per session, *N* = 8 in total), where trials that showed deviation from the fixation were excluded as they did not reflect covert attention.

#### Session One: Behavioural

This first behavioural session included a practice, a staircase and the main behavioural task.

The practice included 20 practice trials with motion coherence set to 99%, followed by A QUEST Staircase (Watson & Pelli, 1979, 1983) designed to adjust the motion coherence of stimuli to achieve 75% accuracy. Two staircases were performed, one for the target appearing in the right hemifield and another for the left. Cue validity during the staircase was 100%. During the main task, the cue-target interval ranged from 600 – 920ms (in steps of 20ms, 6 trials per interval, per block). In total, there were 102 trials in each of the eight blocks of this session, equating to 816 trials in total (*N* = 680 congruent, 136 incongruent). Only participants achieving an overall performance accuracy of >70% on congruent trials and >50% on incongruent trials were invited back to participate in session two (EEG). In addition, performance had to be better for congruent than incongruent trials, ensuring that the participants were covertly shifting their attention based on the cue.

#### Session Two: EEG

After the two staircases, participants performed the main task again, with the addition of extra cue-target intervals of 1000 – 1160ms (steps of 40ms). Session two included 162 trials per block. In total, for this session, there were 1296 trials, of which 816 were behavioural trials and 480 were EEG (*N* = 1080 congruent, 216 incongruent). The 2 sessions yielded 2112 trials per participant (*N* = 1760 congruent, 352 incongruent). Of these, 1632 trials contributed to the behavioural fluctuation analysis, whereas 480 trials (those with cue–target intervals ≥1000ms) entered the EEG analyses. The behavioural fluctuation analysis was restricted to the earlier 600 – 920ms interval to characterise rhythmic changes in performance while minimising contamination from cue-evoked activity, whereas EEG analyses focused on the later interval to isolate sustained pre-target oscillatory dynamics.

EEG data were collected using a 64-channel BioSemi ActiveTwo system (BioSemi, Amsterdam, Netherlands), sampled at 1024 Hz. Three additional electrodes were used, one on each mastoid (right and left) for off-line referencing, and another on the right cheek to measure vertical eye movements (after being referenced to Fp2). Horizontal eye movements were detected by referencing AF7 to AF8 offline.

### Analysis

#### EEG Analysis

All analyses were performed in MATLAB (R2022a, MathWorks, 2022) using custom scripts and the FieldTrip Toolbox (2021 release, https://www.ru.nl/neuroimaging/fieldtrip; Oostenveld et al., 2011). Only trials where the target was presented ≥1000ms after the cue onset were included in the analysis (*N* = 480). The EEG signal was initially epoched from −1000 to 2160ms relative to the cue onset, to include cue and target-related activity. The resulting 480 epochs were referenced to the mastoids, zero-padded to 5 seconds (to avoid filter artifacts), band-pass filtered (1-48 Hz) with a two-pass reverse filter (Gross, 2014) and demeaned to achieve baseline correction. Vertical EOG (VEOG) was calculated to detect blinks and saccades, then subjected to the same pre-processing steps as the EEG signal. Both were combined and resampled to 256 Hz. Eye-tracking (ET) data from each participant was also resampled, imported and concatenated with EEG signals and used to detect microsaccades.

A semi-automatic artifact rejection was used to highlight artifacts (ft_artifact_zvalue, 6 SDs). Each trial was then visually inspected and removed if saccades/microsaccades, movement artifacts, and blinks occurred around the cue or target presentation. On average, approximately 15% of trials were removed (*N* = 406 trials remaining, *SD* = 58.25 trials). Bad channels were also interpolated (*N* = 0.42 channels on average, *SD* = 0.86), and the resulting signal was re-referenced to the average of all channels. Independent component analysis (ICA) was conducted using the *runica* method in FieldTrip to remove remaining eye movements and sustained muscle activity. On average, 5 ICs were identified as artifacts and were removed (*SD* = 2.21). Finally, *ft_rejectvisual* (‘summary’ mode) was used as a concluding quality check.

#### Regression

To estimate the effect of alpha and beta phase on behavioural performance, we looked at the activity before the target presentation. To avoid contamination from post-stimulus activity and filter edge artifacts, we constructed mirrored epochs for each trial. Specifically, the pre-stimulus interval from −500 to −10ms relative to target onset was extracted and then temporally reversed and concatenated to the original segment, creating a symmetric signal around the −10ms boundary. A spectral estimation of the data was then computed for a range of frequencies (7 to 35 Hz, in steps of 1 Hz). This was achieved by first bandpass filtering the raw EEG signals (two-pass reverse) between a range of [*fc* − *hw, fc* + *hw*], where *fc* is the centre frequency and *hw* is frequency width, determining the spectral smoothing. We set hw to be 0.3 times the centre frequency.

Data was passed through a Hilbert Transform to obtain the analytic signal, from which phase angles were extracted for each frequency and time point according to the following formula (1):

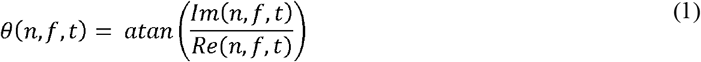

Where *n* is the trial number, *f* is the centre frequency and *t* is the time point. *Im* is the imaginary part of the complex value from Hilbert transformation, whilst *Re* is the real value. After time-frequency decomposition, only the original (non-mirrored) pre-stimulus segment was retained for analysis.

To bring together behavioural and EEG findings, we applied a regression model to determine whether oscillatory phase in the pre-target window can predict *d’* and/or *c* at the single-trial level. Due to the time-consuming nature of this analysis, we selected 51 channels (removing Fp1, Fp2, AF8, Fpz, FT8, FT7, AF7, AF3, AF4, T7, T8, TP7 and TP8) and pre-target timepoints between −250 and −10ms, only looking at alpha and beta frequencies between 7 and 35 Hz. Signals from the two cueing conditions (left, right attentional orienting) were combined, and only congruent trials were selected to remove the confounding effects of attentional reorienting.

Using phase angles, we modelled trial-wise responses using a binomial generalised linear model with a probit link function, following previous approaches to relate oscillatory phase to behaviour (Zoefel et al., 2019; Zhou et al. 2021). Oscillatory phase was entered as circular predictors using its sine and cosine components. These circular predictors were included in our model alongside target movement direction and their combined predictive value (phase angle*movement). For each participant, we modelled the response (signal presence/absence) at each trial according to the following formula (2):

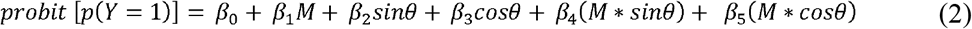

For this, *p*(*Y* = 1) is the probability of answering ‘dots are moving right’ (our signal), *M* is the direction of the target movement (0 = left, 1 = right) and θ is the phase angle. To note, we chose to define dots moving rightward as the signal, in line with the calculations of *d’* and *c* for the behavioural data, although this choice is arbitrary. The output of this model provided beta coefficient values for each of our predictors and their interactions, for every channel, frequency and timepoint. We then utilised the coefficient outputs from the regression analysis to calculate the magnitude of the phase effect on *c* and *d’*. Sensitivity (*d*′; Equation 3.1) is captured by the interaction terms, whereas modulation of the decision criterion (*c*; Equation 3.2) is captured by the main effects of phase. To quantify these effects independently of phase angle, sine and cosine coefficients were combined using a root-mean-square (RMS) metric:

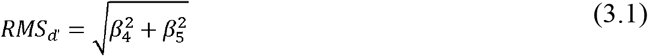

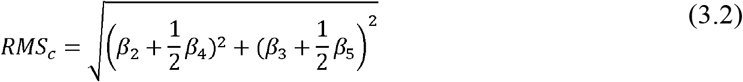

These equations mirror the standard definition of *d’* and *c*. To assess statistical significance, a permutation procedure was implemented at the single-subject level. This analysis was repeated for each participant 100 times for each channel, frequency and timepoint. For each of these iterations, participant responses were randomly shuffled across trials to create a participant-specific null distribution. The number of permutations was limited due to the computational demands of the analysis.

Group level significance was assessed separately for *d’* and *c*, using cluster-based permutation tests across channels, frequencies, and time points (2500 permutations, cluster statistic = *maxsum*). This used a dependent samples *t*-test to determine whether there was a significant difference between the real and permuted distribution of RMS values. Null estimates were obtained by shuffling the trial-wise response vector and recomputing the GLM 100 times, thereby disrupting the relationship between phase and behaviour while preserving the phase and stimulus structure of the data. A final *p*-value was subsequently calculated as the proportion of random samples that resulted in larger *t*-statistics than the real distribution. A one-tailed test was used, as RMS values reflect the magnitude of phase modulation and are strictly non-negative. The critical alpha level was set at *p* = 0.05.

### Multivariate Pattern Analysis

To determine whether pre-target activity could reliably distinguish between trials in which participants shifted their attention to the left versus right, we used artifact-free and low-pass filtered data (40 Hz), as high-pass filtering can bias decoding performance (Van Driel et al., 2021). EEG data were downsampled to 128 Hz to reduce computation time. Data were epoched from −0.5 to 1.5s relative to cue onset to minimise edge artefacts in subsequent time–frequency analyses and baseline-corrected using a −0.2 to 0s window. MVPA was first performed at the single participant level and restricted to pre-target time points. To ensure that decoding reflected successful attentional orienting, only trials with correct responses were included, irrespective of cue validity. As our primary interest lay in visuo-attention processes, and because the regression analysis showed the strongest effects over posterior sensors, classifier performance was examined over a restricted set of occipito-parietal electrodes (Pz, P1, P2, P3, P4, POz, PO3, PO4, Oz, O1 and O2).

We adopted a time generalisation approach, in which a linear discriminant analysis (LDA) classifier was trained at each time point and tested across all other time points of the selected EEG signal. This resulted in 2D matrix of classification performance across all train × test time combinations (Treder,D2020). We implemented this analysis in MATLAB using the MVPA-Light toolbox (Treder, 2020; https://github.com/treder/MVPA-Light.git). Cross-validation was performed using predefined, block-based folds, constructed such that each fold contained trials from both attentional shift conditions (one left and one right block per fold), while ensuring independence between training and test sets. Classifier performance was quantified using the area under the receiver operating characteristic curve (AUC), chosen over accuracy as it provides a more robust metric in the presence of potential class imbalance.

Statistical significance of decoding performance was assessed using a non-parametric cluster-based permutation framework. Statistical testing was restricted to the post-cue window (0 – 1s). At the single-participant level, null distributions were generated by repeating the classification procedure with labels permuted at the block level within each fold, preserving the temporal and block structure of the data while disrupting the relationship between neural activity and attentional shift direction (50 permutations per participant). At the group level, subject-level decoding matrices were compared against chance performance (AUC = 0.5) using cluster-based permutation testing across the full train × test time matrix.

To characterise the spectral composition of decoding-relevant activity, each participant’s time generalisation matrix was first expressed relative to chance (AUC − 0.5), detrended along both training and test dimensions, and mean-centred across rows and columns. Time-frequency decomposition was performed using a sliding-window convolution approach (*mtmconvol* in Fieldtrip, Hanning taper), with time windows corresponding to three cycles per frequency (7 – 35 Hz). Decomposition was performed separately along the training and test dimensions, and the resulting power estimates were averaged to yield a single time-frequency representation per train × test time point. Finally, the time-window was again restricted to the post-cue window.

Group-level significance of spectral power was evaluated using a permutation-based null distribution. At the single-participant level, 50 label-shuffled classifiers were computed, and at the group level, null distributions were constructed by randomly sampling one permutation per participant across 10,000 iterations. Empirical *p*-values were computed for each train x test x frequency voxel and corrected for multiple comparisons using false discovery rate (FDR) procedures. Importantly, time– frequency statistics were evaluated only within train–test time points that showed significant decoding, ensuring that spectral effects were interpreted strictly in the context of successful classification.

For visualisation, we constructed a frequency map across the train × test time plane. For each voxel showing both significant decoding and at least one frequency surviving FDR correction, we assigned the frequency with the largest significant power. Crucially, this procedure identifies the dominant frequency at each decoding-relevant voxel and does not imply the absence of other oscillatory components. Frequencies that did not exceed the statistical threshold, or that were weaker than the dominant component, were not represented in this map. Voxels without any significant frequency were left undefined. Consequently, these maps highlight the frequency band contributing most strongly to reliable decoding at each time point, rather than providing an exhaustive representation of all underlying oscillatory activity present in the data.

### Behavioural Analysis

#### Preliminary checks

Behavioural data obtained from both sessions were combined and analysed together. Trials containing blinks or saccades occurring between 100ms before and 60ms after target onset were excluded based on eye-tracking data. Following these exclusions, an average of 1505 trials per participant remained for analysis. Overall task performance was assessed using accuracy, perceptual sensitivity (*d*′), and decision criterion (*c*). These measures were calculated separately for congruent and incongruent trials and compared using paired-samples *t*-tests to verify that participants benefited from the spatial cue and were appropriately allocating attention. Additional analyses confirmed that performance did not differ across sessions (see Document S1).

#### Main analysis

Behavioural data from both sessions were combined, and *d*′ and *c* were computed separately for each cue-to-target interval using only congruent trials, thereby minimising the contribution of attentional reorienting elicited by invalid cues. The following equations were used:

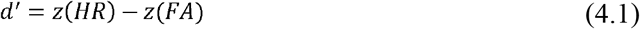

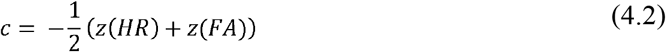

Where HR and FA are the hit rate and false alarm rate, respectively, and *z*(*x*) is the inverse of the normal cumulative function evaluated at *x* (Macmillan, 2002; Macmillan & Creelman, 1991). Rightward motion was arbitrarily defined as the signal, and leftward motion as noise. This yielded a time course of visual performance covering a cue-to-target interval from 600 to 920ms (steps of 20ms) post cue. This was subsequently analysed for the presence of cyclic patterns (see below).

Additionally, as behavioural and EEG data covered slightly different windows (i.e., 600-920ms vs 100-1160ms), we also verified that performance did not differ across windows (Document S1).

### Fluctuation in Behaviour

Both *d’* and *c* time course data were linearly detrended to remove the linear effects across inter-stimulus intervals and retain any cyclic patterns around the mean. We then tested for the presence of cyclic patterns by employing a curve-fitting procedure using robust nonlinear least-squares fitting. For each participant, we fitted cosine models C(t) between 7 and 25 Hz (Nyquist frequency) in 1 Hz steps according to the following formula:

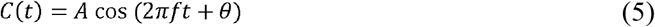

Where *A, f* and *θ* are the amplitude, frequency, and phase of the cosine, respectively.

Resulting R-squared values for each frequency model were then extracted and tested for significance using a bootstrap procedure. Cue-to-target interval labels were randomly permuted over 500 iterations, with cosine models fitted to the resulting behavioural pattern of each participant, generating a null distribution of 500 R-squared values. The R-squared values obtained from the actual data were compared to the null distribution and considered to be significant if they fell in the top 97.5th percentile.

To verify that significant behavioural fluctuations reflected genuine oscillatory activity rather than aperiodic spectral structure, irregular resampling auto-spectral analysis (IRASA; Wen and Liu, 2016) was subsequently applied to the *d*′ and *c* time courses. IRASA separates oscillatory and fractal (1/*f*) components of the power spectrum, enabling rhythmic fluctuations to be distinguished from broadband spectral contributions. Significant frequencies identified by the cosine-fitting procedure were interpreted as oscillatory only if they were also evident in the IRASA-derived oscillatory component, rather than being explained by the fractal component (Helfrich et al., 2018).

## Results

We first confirmed that participants shifted their attention in line with the block instruction. Performance remained at ~75% accuracy during congruent trials (*M* = 74.26, *SE* = 1.12) and above chance during incongruent trials (*M* = 60.51, *SE* = 1.08; Figure 1B), thus showing an attentional benefit (*t*(25) = 8.9, *p* < .001, *d* = 1.78). We additionally confirmed this attentional shift on a neural level, where leftward versus rightward covert attention shifts displayed the classic alpha-signature of attentional orienting over the occipital electrodes (Figure 1C). This was characterised by a significant decrease of alpha power over the left (negative cluster, *p* = .018) and an increase over the right (positive cluster, *p* = .036) occipital electrodes, confirming that participants validly utilised the cue to shift their attention for the upcoming target onset. The alpha signature was not identified at pre-cue time points (negative cluster, *p* = .22), confirming that the participants used the cue to initiate a covert attentional shift as intended.

### Beta phase tracks decision criterion and perceptual sensitivity

Inspired by recent findings linking beta phase to perceptual performance and sensory excitability, we asked whether the ongoing oscillatory phase predicts perceptual sensitivity (*d*′) and/or decision criterion (*c*). Although we predicted a beta dominance, we also included alpha activity within this analysis, given its known involvement in visuospatial attention.

We estimated instantaneous pre-target phase (7–35 Hz) and used a probit GLM to model trial-wise responses (“rightward” reports) as a function of sine and cosine components of phase, motion direction (left vs right), and their interaction. This enabled the separation of effects, where the phase-dependent modulation of sensitivity is captured by the interaction between phase and stimulus direction, whereas shifts in criterion are reflected in the main effect of phase (see Methods for details).

Our results showed that pre-target beta phase (20–30 Hz) significantly modulated both decision criterion and perceptual sensitivity (Figure 2). However, effects on *c* emerged earlier, between 225 – 175ms before target onset, and were distributed across fronto-central and occipital electrodes (*p* = .002; Figure 2A). Effects on *d’* were stronger over posterior channels and occurred closer to target onset, between 150 – 100ms before the target onset (*p* = .0008; Figure 2B). Taken together, these findings demonstrate that pre-target beta phase is associated with distinct behavioural components of perceptual decision-making.

**Figure 2.**
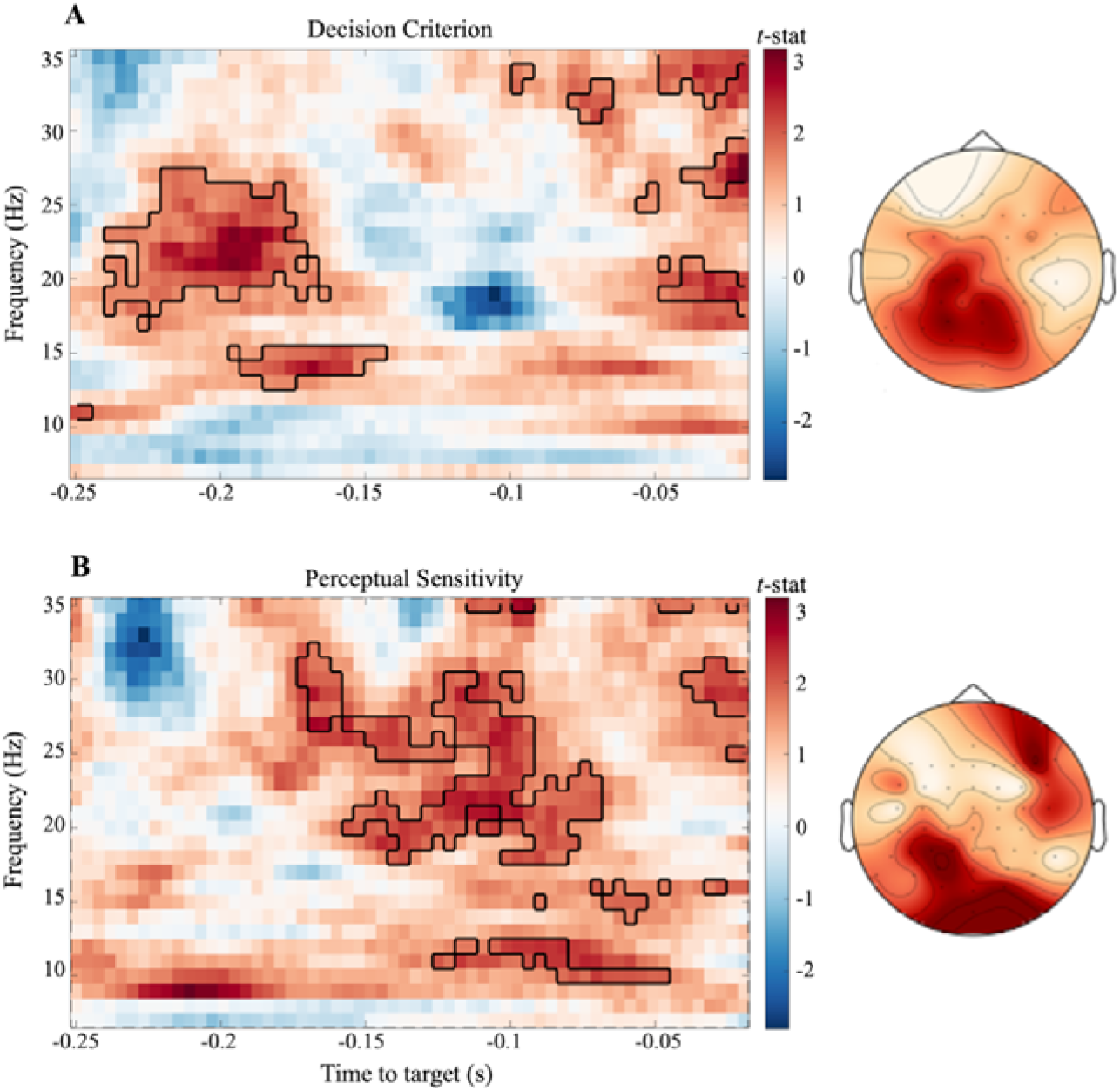
Pre-target beta phase predicts decision criterion and perceptual sensitivity. Time-frequency representations illustrating the strength of phase-behaviour coupling derived from the generalised linear model (GLM). Panels show effects of oscillatory phase on A) decision criterion (*c*; main effect of phase) and B) perceptual sensitivity (*d*_′_; phase × stimulus interaction). Colours represent group-level test statistics of the root-mean-square (RMS) magnitude of phase modulation, obtained by combining sine and cosine phase coefficients independently of preferred phase angle. Black contours denote clusters surviving cluster-based permutation testing across channels, frequencies, and time points. For visualisation, time-frequency maps are shown at electrode Oz; however, statistical inference was performed across the full sensor array. Right panels display the spatial topographies of the corresponding significant clusters. The x-axis indicates time relative to target onset (zero).

### Beta-band signatures in behavioural metrics

We next examined whether behavioural performance itself showed evidence of rhythmic modulation (*c* and *d’*; Figure 3A and S3A, respectively) by sampling behaviour at earlier post□cue delays than those entering the EEG analysis. This created a measure of oscillations without the caveat of ERP influences caused by the cue presentation. Both *d’* and *c* time series were analysed for the presence of cyclic patterns by fitting cosine models (7 and 25 Hz), with the addition of IRASA estimating the background 1/*f* spectrum (see Figure 3C and S3C).

**Figure 3.**
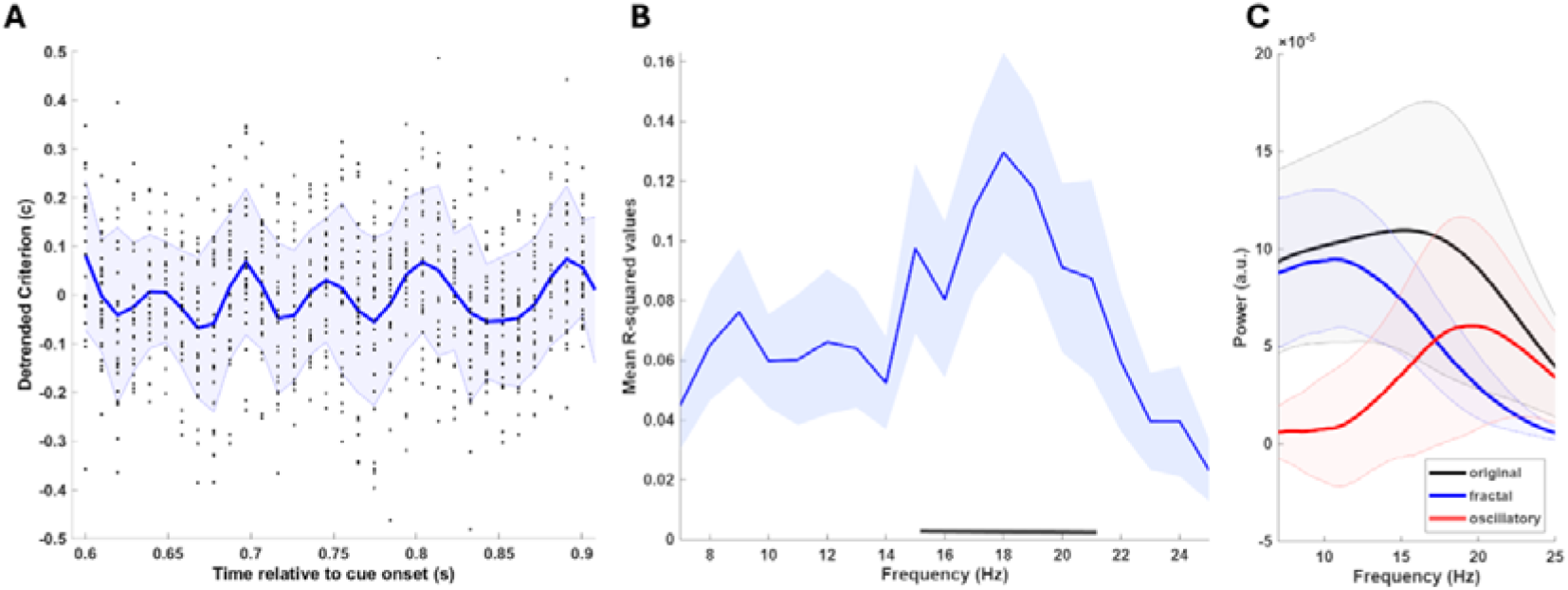
Rhythmic fluctuation in behavioural measures. A) Time courses of the decision criterion (*c*) plotted for individual participants (dots) and the group average (blue line). Shaded regions indicate ±1 SD. The x-axis denotes time relative to cue onset. B) R-squared values plotted as a function of fitted cosine frequency. The black line indicates models significantly fitting the data; fits exceeding the 97.5th percentile of the null distribution were considered statistically significant. C) Separation of aperiodic (1/*f*) and oscillatory components for measures of *c*. The original spectrum is shown in black, the estimated aperiodic component in blue, and the oscillatory component in red. Only oscillatory activity exceeding the aperiodic estimate was considered meaningful. Lines represent the group average, and shaded areas indicate ±1 SD. Significant model fits appeared at ~15–21 Hz, exceeding both the permutation threshold and the aperiodic (1/*f*) estimate.

Across participants, *c* fluctuated over time and in line with the EEG results, the frequency best explaining this measure pointed to beta-band oscillations. R-squared values within the beta range showed a significant fit for models of ~15 – 21 Hz (Figure 3B), which was confirmed to be above the aperiodic activity estimation. For *d’*, the bootstrapping procedure highlighted several significant models (Figure S3B), which, however, were not confirmed by the IRASA decomposition (Figure S3C).

Taken together, the behavioural data converge with EEG findings in suggesting that beta□band activity in an early post□cue time window is associated with systematic changes in the decision criterion, fluctuating with a beta oscillation.

### Attentional orienting is preferentially decoded from alpha-band activity

We next asked whether attentional orienting could be decoded from pre-target EEG activity. To this end, we trained a time-resolved classifier to distinguish leftward from rightward attentional shifts and generated a time-generalisation matrix (TGM) for each participant (Figure 4A). Non-parametric cluster-based permutation revealed an above-chance performance emerging 200ms post-cue (peak AUC = 0.61 ± 0.03 SD across participants and significant window; Figure 4B). Inspection of the temporal generalisation matrices revealed that decoding was not confined to the diagonal but instead exhibited broad off-diagonal generalisation. Specifically, classifiers trained at later time points (~300ms post-cue) generalised across an extended range of subsequent time points, forming a large, sustained cluster of above-chance decoding. This pattern indicates that attentional orienting is supported by a temporally stable neural representation, rather than by rapidly changing, transient cue-locked processes. Importantly, ERP analyses revealed no significant differences between leftward and rightward attentional orienting conditions (see Figure S4), further supporting that decoding performance was unlikely to be driven by condition-specific evoked responses.

**Figure 4.**
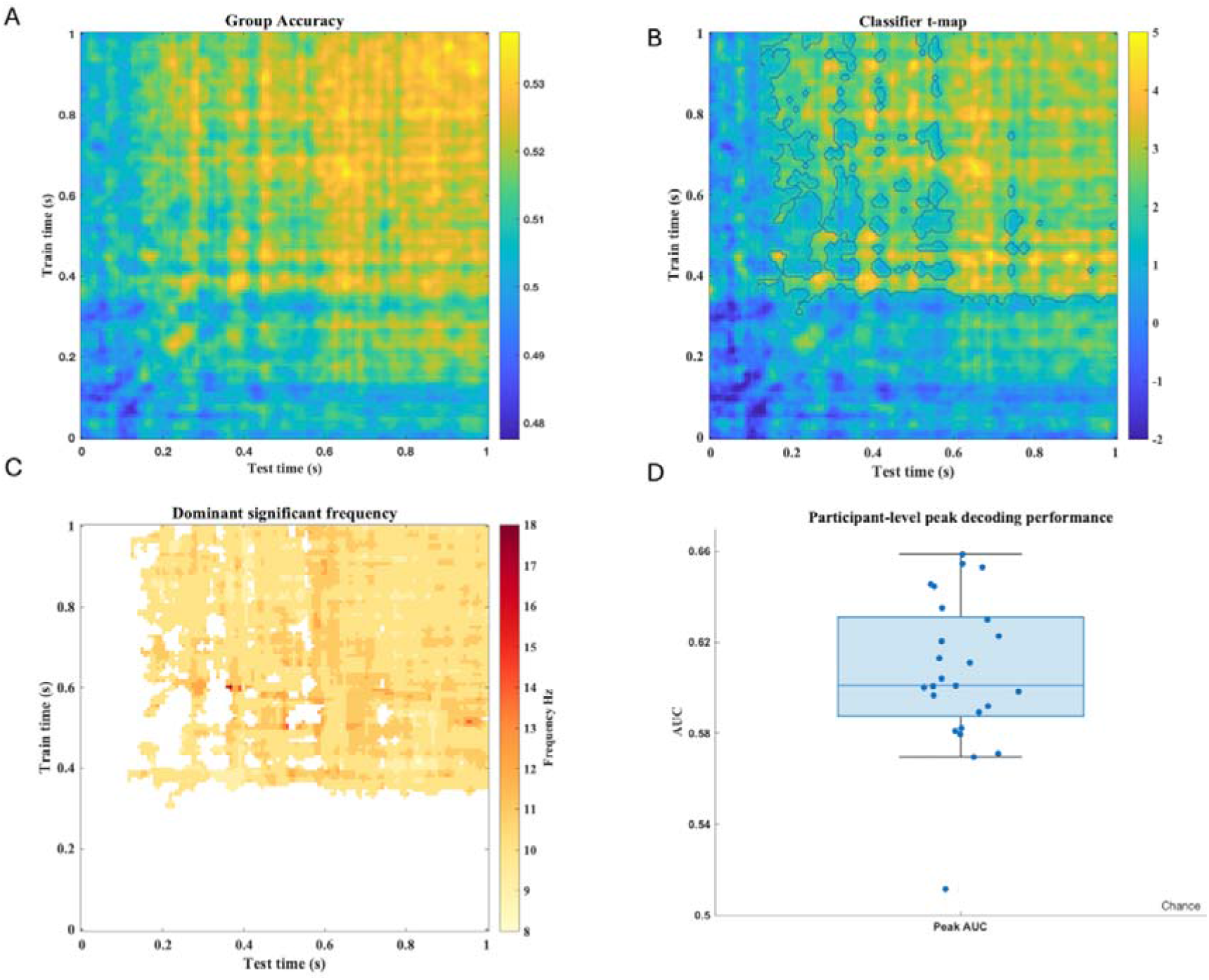
Decoding the direction of attentional orienting from pre-target EEG activity. A) Group-averaged time-generalisation matrix showing classifier performance (area under the curve, AUC) for decoding left versus right attentional shifts from occipito-parietal EEG signals. Warmer colours indicate above-chance decoding performance, cooler colours indicate lower performance. B) Statistical map of decoding performance, with contours outlining clusters that survived non-parametric cluster-based permutation testing against chance. C) Dominant oscillatory frequency associated with decoding performance, extracted from train × test time points showing significant classifier performance. Colours represent the frequency (in Hz) at which time-frequency power was maximal within each decoding-significant voxel. D) Participant-level peak decoding performance (AUC) across subjects. The boxplot depicts the median (line), the interquartile range (box), 1.5 times the interquartile range (whiskers).

To identify the oscillatory components contributing to this decoding effect, we performed time-frequency analyses on the TGM of each participant using the time window that exhibited significant decoding performance (as in Melcón et al., 2025). Frequencies between 8 – 18 Hz survived statistical thresholding; however, as clearly shown in Figure 4C, fluctuations in decoding accuracy were mostly driven by alpha-band activity. Together, these findings indicate that alpha-band dynamics contributed most strongly to decoding attentional orienting states (see Figure S2A for beta results).

## Discussion

The present study set out to characterise how visuospatial attention shapes perceptual decisions through oscillatory dynamics in the human visual system. By combining a sustained spatial cueing paradigm with signal detection theory measures and time-resolved EEG analyses, we demonstrate a functional dissociation between alpha- and beta-band dynamics: occipital beta phase predicted behavioural outcomes, whereas alpha frequency dominated the neural discriminability of attentional orienting states. Specifically, we show that beta phase can predict decision criterion (*c*) at earlier pre-target intervals and perceptual sensitivity (*d’*) at later intervals, closer to target onset. This suggests that occipital beta oscillations contribute dynamically to different aspects of attentional processing, extending previous work that links beta phase to discrimination accuracy, phosphene detection and visual cortex excitability (Samaha, Gosseries, et al., 2017; Trajkovic et al., 2025; Veniero et al., 2021).

Our findings further suggest that occipital beta phase is not just related to sensory gain but also influences the decisional processes. Specifically, beta phase was associated with participants’ decision-related response bias (*c*) and how effectively they extract target sensory information (*d’*), with each effect depending on temporal proximity to the upcoming stimulus. Of note, the early involvement of beta in relation to the decision criterion was demonstrated by the EEG data and supported by its behavioural fluctuation. This extends previous findings, showing that beta-band oscillations reflect the temporal and spatial dynamics underlying the accumulation and processing of task-relevant information within the sensorimotor network, ultimately shaping decision outcomes (Haegens et al., 2011). Our results build on these findings by demonstrating a temporal progression in human observers, whereby beta phase predicted criterion earlier than sensitivity, suggesting partially dissociable stages of perceptual decision formation (Haegens et al., 2011). Earlier and later beta-phase associations may therefore relate to distinct aspects of response bias and perceptual sensitivity at different pre-target intervals.

Previous work examining oscillatory influences on *d’* and *c* has produced mixed findings and has largely focused on perceptual rather than attentional processes (Chanes et al., 2013; Ho et al., 2017; Iemi et al., 2017; Limbach & Corballis, 2016; Samaha et al., 2020; Samaha, Iemi, et al., 2017; Sherman et al., 2016; Zhou et al., 2021). Much of this literature has also specifically focused on alpha-band dynamics. For example, some studies report that alpha phase predicts criterion but not sensitivity (Sherman et al., 2016), whereas others show that fluctuations in alpha power modulate sensitivity but not criterion (Zhou et al., 2021). Conversely, several investigations find the opposite pattern, with reduced alpha power biasing participants toward reporting stimulus presence, reflecting shifts of *c* rather than *d*′ (Iemi et al., 2017; Samaha et al., 2020; Samaha, Iemi, et al., 2017). These inconsistencies may stem from differences in task demands or analytical approaches, or they may indicate that alpha dynamics do not map cleanly onto a single SDT measure. Importantly, however, these studies were primarily designed to link ongoing oscillatory activity, typically alpha, to visual perception and characteristically did not require endogenous shifts of visuospatial attention. Thus, although previous work implicates oscillatory activity in shaping perceptual decisions, it has remained unclear whether the same mechanisms operate during top-down attentional orienting.

Our findings, therefore, contribute new evidence by demonstrating that when attentional orienting is involved, there is an accompanying oscillatory modulation of *d*′ and *c*, specifically within an attentional context. This dissociation between perceptual sensitivity and decision criterion is broadly consistent with frequency-specific TMS studies showing that beta-frequency stimulation enhances perceptual sensitivity, whereas higher stimulation frequencies preferentially modulate criterion (Chanes et al., 2013). The predominance of beta-band effects in our study, compared with the alpha-driven findings of perceptual research, likely reflects the fundamentally different demands of endogenous attentional orienting. Beta-band activity has, in fact, been associated with top-down mechanisms, with fluctuations in this frequency range cyclically modulating the visibility of target stimuli within attentionally prioritised regions (Cabeza, 2002; Richter et al., 2017; Veniero et al., 2021). Importantly, neuromodulation studies have shown that top-down attentional signals generated through stimulation of the frontal eye fields (FEFs) induce a phase reset of beta-band activity in posterior-occipital regions, shaping the rhythmic sampling of visual stimuli (Trajkovic et al., 2025; Veniero et al., 2021). This aligns with hierarchical models of cortical communication, in which beta-band activity is thought to mediate long-range feedback from higher-order areas to sensory cortices, transmitting top-down signals and mediating attentional processes (Bastos et al., 2015; Michalareas et al., 2016; Spitzer & Haegens, 2017). Consistent with this view, synchronisation along the dorsal visual pathway has been shown to track the current focus of spatial attention (Di Dona & Ronconi, 2023; Siegel et al., 2008). This provides a context for interpreting the beta-phase effects observed here, consistent with the possibility that beta oscillations contribute to the temporal organisation of perceptual sampling alongside complementary oscillatory mechanisms of visuospatial attention, including alpha. However, as we did not directly measure alpha–beta interactions, this interpretation remains speculative.

Our second main finding concerns the neural discriminability of attentional orienting states. Decoding performance was dominated by alpha-band activity, indicating that alpha dynamics contributed most strongly to distinguishing leftward from rightward attentional orienting. Importantly, decoding generalised across non-identical time points, suggesting that attentional-state information was maintained across sustained pre-target intervals rather than being confined to transient cue-locked responses. These findings are consistent with frameworks proposing that attentional orienting involves dynamically fluctuating neural states, potentially linked to rhythmic attentional sampling mechanisms (Helfrich et al., 2018; Landau & Fries, 2012).

Although speculative, our findings also align closely with the idea that periods of time dedicated to stimulus processing are characterised by beta-band activity associated with attentional stability and suppression of attentional shifts, whereas alpha-band activity reflects transient states of sensory attenuation and increased susceptibility to attentional reorienting (Fiebelkorn & Kastner, 2019). Specifically, our finding that beta phase predicted both decision criterion and perceptual sensitivity suggests that beta-band dynamics contribute to the temporal organisation of attentional stability and perceptual decision formation. By contrast, alpha-frequency structure dominated the rhythmic discriminability of attended location, consistent with the idea that alpha-band dynamics regulate the periodic gating of sensory information during sustained attentional orienting.

More broadly, our results suggest that rhythmic attentional sampling may emerge through potentially complementary contributions of beta- and alpha-band dynamics. In conclusion, these findings support a functional dissociation between alpha and beta rhythms in visuospatial attention. Such division of labour aligns with emerging frameworks proposing that beta oscillations coordinate top-down signals of attention (Richter et al., 2017; Riddle et al., 2019), whilst alpha regulates sensory sampling (Thut et al., 2006). In clearer terms, alpha indexes *where* attention is allocated, and beta shapes *how* and *how well* perceptual decisions are made.

