## SupplementaryMaterials for "Alpha oscillations support attentional orienting while beta supports perceptual decision-making"

**Document S1.**

When comparing performance across sessions, accuracy did not differ between session one (*M* = 67.52, *SE* = .8) and two (*M* = 67.25, *SE* = .99), *t*(25) = .3, *p* = .38. Similarly, there were no session differences for *d’* (session one: *M* = .97, *SE* = .047; session two: *M* = .96, *SE* = .057), *t*(25) = .093, *p* = .46, or for *c* (session one: *M* = .-13, *SE* = .041; session two: *M* = -.095, *SE* = .051), *t*(25) = -.95, *p* = .18.

In contrast, when merging data from both sessions, performance did significantly differ between congruent and incongruent trials across all three behavioural measures. As expected, accuracy was significantly higher for congruent trials (*M* = 74.05, *SE* = .01) than incongruent (*M* = 60.45, *SE* = .02), *t*(25) = 7.50, *p* < .001, *d* = 1.47, 95% CI [.91, 1.96]. This was the same pattern of results for *d’* (congruent trials: *M* = 1.35, *SE* = .074; incongruent: *M* = .57, *SE* = .08), *t*(25) = 7.58, *p* < .001, *d* = 1.49, 95% CI [.92, 2.04], but not for *c* (congruent trials: *M* = -.08, *SE* = .047; incongruent: *M* = -.11, *SE* = .06), *t*(25) = .82, *p* = .42.

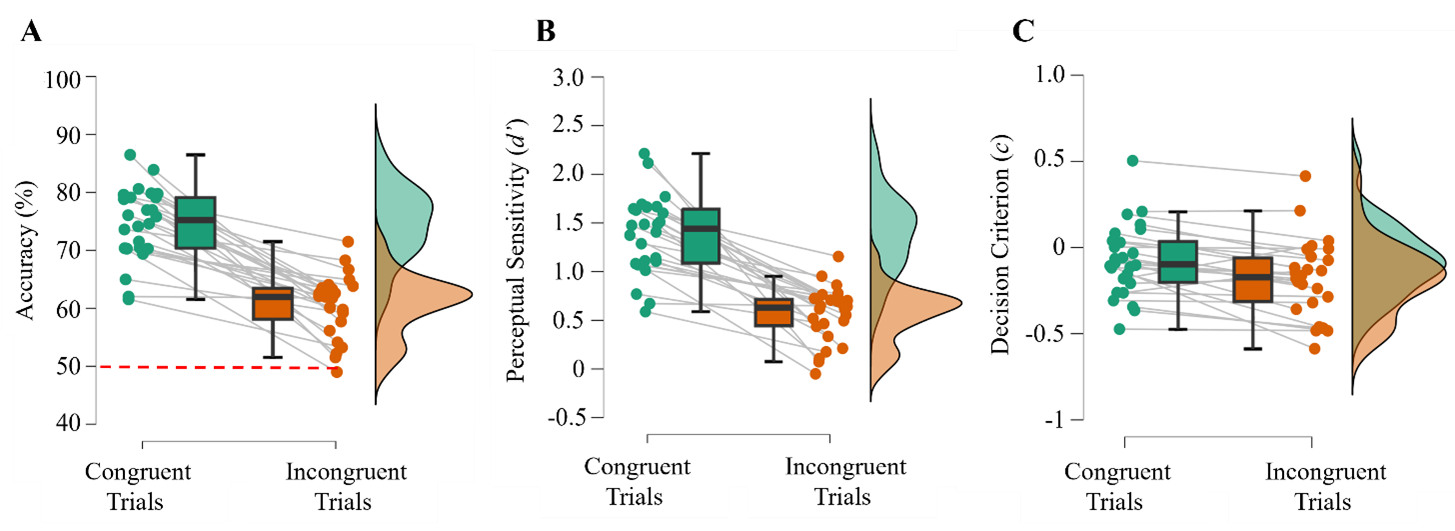

***Figure* S1**. A) Performance accuracy for congruent and incongruent trials. Individual participants are shown as dots; box midlines represent the mean. Chance‑level performance (50%) is indicated by a dashed red line. B) Perceptual sensitivity (*d′*) and C. decision criterion (*c*) are plotted for congruent and incongruent trials using the same format.

To additionally verify that the performance in the behavioural (latency: 600 – 920ms from cue) and EEG window (latency: 1000 – 1160ms from the cue) was comparable, we conducted a *t*-test to determine that on average, performance was near identical for both *d’* (window 1: *M* = 1.37, *SD* = .40, window 2: *M* = 1.35, *SD* = .38) and *c* (window 1: *M* = -.075, *SD* = .21, window 2: *M* = -.08, *SD* = .24). A *t*-test confirmed no statistically significant difference across the two for *d’* (*t*(25) = .38, *p* = .71) and *c* (*t*(25) = .21, *p* = .84). This suggested that the two windows contained the same signals and so we can extrapolate findings from one window into the other.

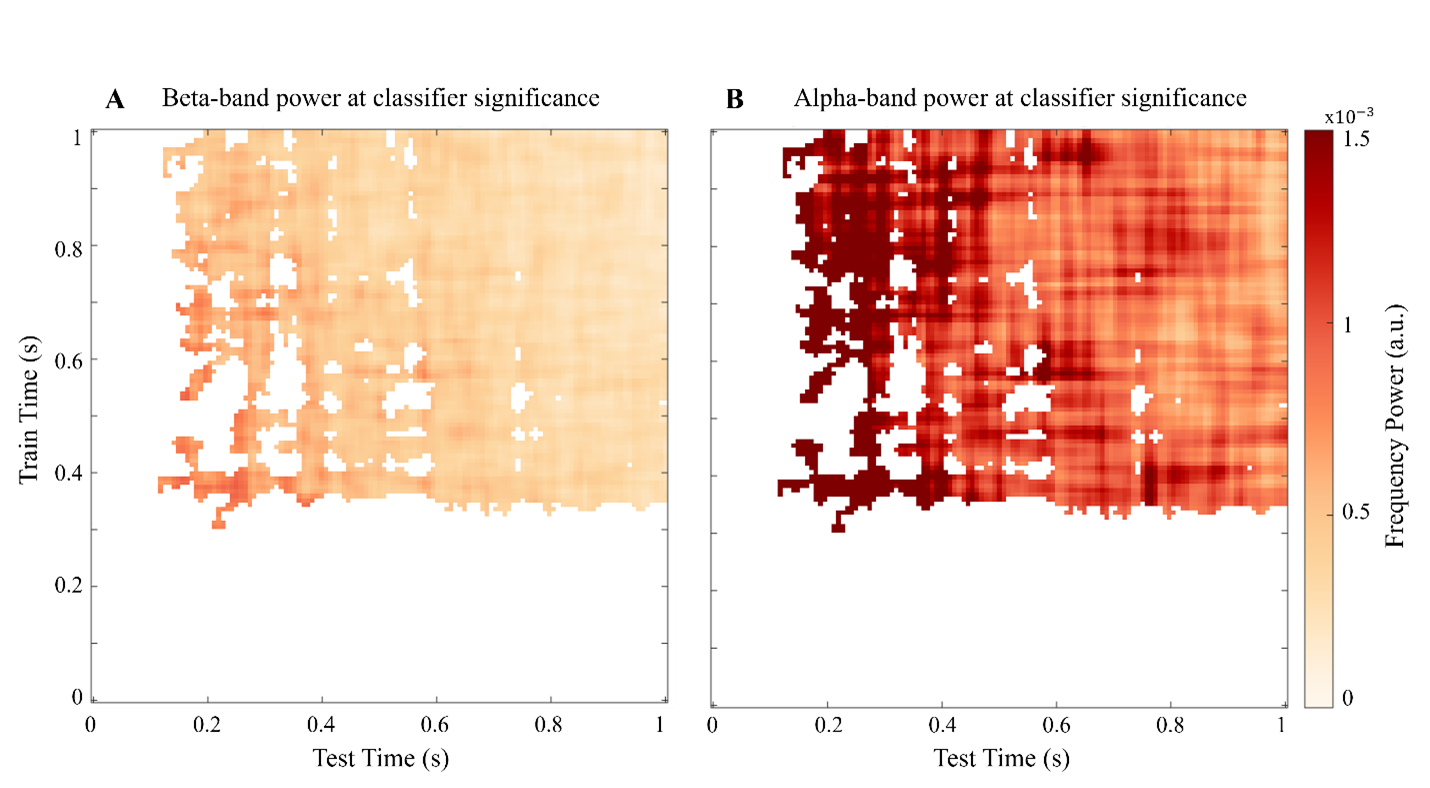
***Figure* S2.**

***Figure* S2**. Frequency‑specific contributions to classifier performance at significant time points. A) Beta‑band power contributions and B) alpha‑band power contributions. Frequency power (a.u.) is illustrated in the colour map, with lighter orange indicating lower power and deeper red indicating higher. Here, alpha power is most dominant in modulating classifier performance.

***Figure* S3.**

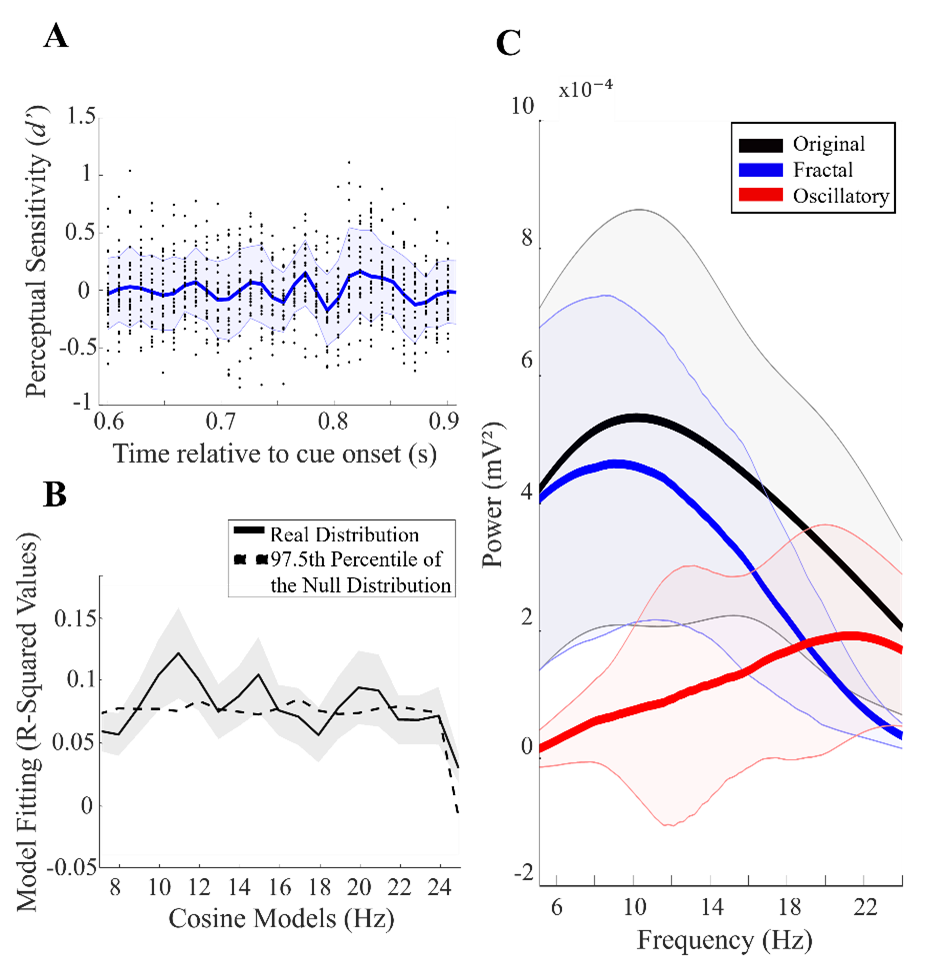

***Figure* S3**. A) Time courses of perceptual sensitivity (*d’*) plotted for individual participants (dots) and the group average (blue line). Shaded regions indicate ±1 SD. The x-axis denotes time relative to cue onset. B) Following model fitting, R-squared values are also plotted from cosine model fits for *d’*. The dashed black line indicates the 97.5th percentile of the null distribution; fits exceeding this threshold were considered statistically significant. C) Separation of aperiodic (1/*f*) and oscillatory components for measures of *d’*. The original spectrum is shown in black, the estimated aperiodic component in blue, and the oscillatory component in red. Only oscillatory activity exceeding the aperiodic estimate was considered meaningful. Lines represent the group average, and shaded areas indicate ±1 SD. For *d′*, cosine model fits exceeded the bootstrap significance threshold but did not exceed the aperiodic (1/*f*) and were therefore not interpreted as reflecting robust oscillatory activity.

***Figure* S4.**

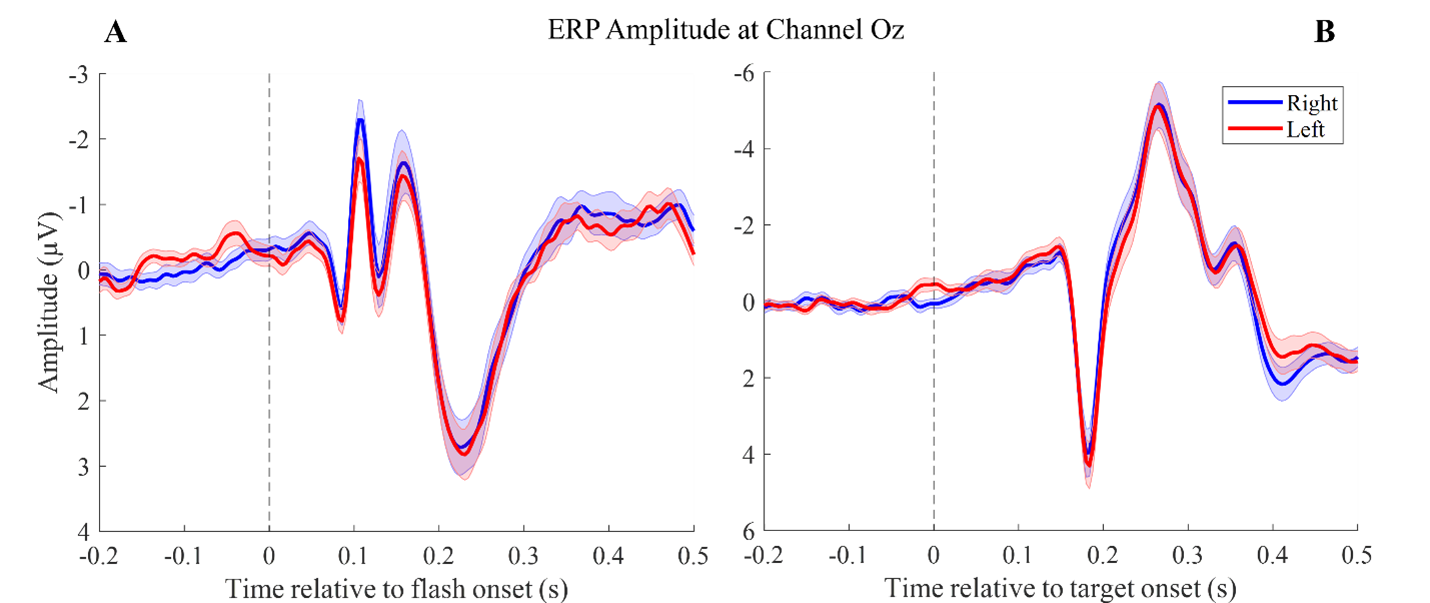

***Figure* S4**. Group‑averaged ERPs for shift‑right (blue) and shift‑left (red) trials, time‑locked to A) cue onset and B) target onset. Signals are shown from channel Oz to illustrate occipital activity without confounds from lateralised responses
